# Ambulatory electrocardiographic longitudinal monitoring in dystrophin-deficient dogs identifies decreased Very Low Frequency power as a hallmark of impaired Heart Rate Variability in Duchenne Muscular Dystrophy

**DOI:** 10.1101/2023.05.08.539882

**Authors:** Inès Barthélémy, Jin Bo Su, Xavier Cauchois, Frédéric Relaix, Bijan Ghaleh, Stéphane Blot

## Abstract

**Background:** Duchenne muscular dystrophy (DMD) patients exhibit a late left ventricular systolic dysfunction preceded by an occult phase, during which myocardial fibrosis progresses and some early functional impairments can be detected. These latter include electrocardiographic (ECG) and heart rate variability (HRV) abnormalities.

**Methods:** A longitudinal study aiming at describing the sequence of ECG and HRV abnormalities, relative to cardiac function degradation, using Holter ECG, was performed in the GRMD (Golden retriever muscular dystrophy) dog model, known to develop a DMD-like disease, including cardiomyopathy.

**Results:** Most of the known ECG and HRV abnormalities described in DMD patients were also found in GRMD dogs, and several of them could be detected months before the decrease of fractional shortening. One of the earliest evidenced abnormalities was a decrease in the very low frequency (VLF) component of the power spectrum, and this decrease was correlated with the further reduction of fractional shortening. Such decreased VLF probably reflects impaired autonomic function and abnormal vasomotor tone.

**Conclusion:** This study provides new insights into the knowledge of the GRMD dog model and DMD cardiomyopathy and emphasizes the interest to monitor the VLF power in DMD patients, still unexplored in this disease, whilst it is highly predictive of deleterious clinical events in many other pathological conditions.

**Abbreviations:**

cTpnI: Cardiac Troponin I
DMD: Duchenne Muscular Dystrophy
ECG: Electrocardiogram
GRMD: Golden Retriever Muscular Dystrophy
HF: High Frequency
HR: Heart Rate
HRV: Heart Rate Variability
LF: Low Frequency
LGE: Late Gadolinium Enhancement
LTV: Long-Term Variability
LVEF: Left Ventricular Ejection Fraction
NT-proBNP: N-Terminal part of the pro-Brain Natriuretic Peptide
pNN50: percentage of interval differences of successive RR intervals of more than 50 ms
pNN10 %(meanRR): percentage of interval differences of successive RR intervals of more than 10% of the mean RR
QTc: corrected QT interval
RMSSD: square root of the mean squared differences of successive RR intervals
SDNN: Standard Deviation of the RR intervals
LVFS: Left Ventricular Fractional Shortening
STV: Short-Term Variability
VLF: Very Low Frequency
PVB: Premature Ventricular Beat
VT: Ventricular Tachycardia.

## Introduction

Duchenne Muscular Dystrophy (DMD) is an X-linked genetic disorder affecting 15.9 to 19.6 boys over 100000 male births and is due to mutations in the dystrophin gene (1). In skeletal muscles and heart, dystrophin is localized beneath the sarcolemma and stabilizes the cell membrane during contraction. In the absence of dystrophin, the membrane becomes damaged upon mechanical stress and a cascade of events leads to cell degeneration (2). Clinically, DMD is characterized by prominent muscle weakness leading to locomotor disabilities, progressing in the childhood. Complete loss of ambulation occurs around 10 years of age without any therapeutic intervention (1). In the second decade, respiratory muscle weakness progresses, accelerated by the transition to wheelchair, and necessitates respiratory assistance, that is adapted in function of the severity of the respiratory insufficiency (3). Thanks to the progresses made in the medical management of DMD patients over the last decades, in particular in ventilatory assistance, survival of patients has increased by a decade, making the DMD cardiomyopathy a growing medical issue (4). Indeed, clinical cardiomyopathy in DMD appears later than skeletal muscle disabilities, usually in the mid-teens, *i.e*., close to the life expectancy of DMD patients before the addition of ventilatory support (5). With the improvement of survival, cardiomyopathy is becoming an important cause of death in DMD patients and a major therapeutic target (6).

Therefore, standards of care for DMD patients now include a yearly comprehensive cardiac evaluation from time of diagnosis, notably echocardiography, in order to detect the onset of left ventricular (LV) ejection fraction (LVEF) or fractional shortening (LVFS) decrease, and adapt the therapeutic management (3). It is however now well known that before the LVEF decrease, the dystrophin-deficient heart undergoes an “occult” phase during which some early abnormalities can be detected and could help anticipate the drop of LVEF. It is the case of myocardial fibrosis that can be detected months before LV global systolic dysfunction (*i.e*., decrease of the LVEF or LVFS) using cardiac MRI and late gadolinium enhancement (LGE) mapping. Some studies show that the LGE foci and their distribution can be useful in grading the advancement of the cardiac disease, and predict the decrease of LVEF (7, 8). Cardiac MRI is therefore the imaging method that is recommended for the follow-up of DMD patients (3). However, it is unfortunately not accessible to all of them. Some other studies have shown that early echocardiographic abnormalities can be evidenced in young DMD patients with a normal LVEF. Doppler tissue imaging (DTI) techniques allows the identification of an early decrease of radial strain rate (endo-epicardial gradient of velocity) and peak systolic strain of the LV free wall (9). Electrocardiograms (ECG) can also detect abnormalities long before the decrease of LVEF, and yearly ECGs are part of the standards of care for DMD patients (3, 10, 11). Among typical ECG abnormalities, PR shortening, prolonged QT, increased heart rate are found, as well as deep Q-waves and tall R waves on some leads (5, 10, 11). Fragmented QRS can also be found, and are associated with the degree of LV dysfunction and myocardial fibrosis (12). A special attention is paid to the presence of arrhythmic events on ECGs from DMD patients. These arrhythmias can be of supraventricular or ventricular origin, and ventricular premature contractions can lead to ventricular tachycardia episodes or even fibrillation and increase the risk of sudden death (13). Holter electrocardiograms are recommended to detect such arrhythmic events (3). During the early occult phase of cardiomyopathy, heart rate variability (HRV) abnormalities can be detected on ECGs of DMD patients. The studies that focused on this aspect used mostly 24 h Holter ECGs and showed a decreased HRV based on time-domain indices, a decreased high-frequency power and an increased ratio of low-frequency on high-frequency powers using frequency-domain analysis, together with a modified circadian variability (14–20). All these abnormalities evidence an autonomic dysfunction, with increased sympathetic and decreased parasympathetic modulation, that occurs before LV global systolic dysfunction, together with myocardial fibrosis, and aggravates with advanced stages of the disease. HRV decrease has been associated with many other conditions as a prognostic marker of a poor outcome and mortality (21, 22). It has been suggested to use HRV analysis more widely in DMD patients, to improve the early detection of the cardiac disease. However, the sequential evolution of all the early abnormalities that ultimately lead to impaired heart function and the way they interact together in DMD are not yet fully characterized.

To better understand the human DMD, and assess therapeutic strategies, animal models can be useful to provide relevant information. In the field of DMD cardiomyopathy, the canine GRMD (Golden retriever muscular dystrophy) model is noticeably relevant. Indeed, these dogs develop a cardiomyopathy that resembles the human one in many aspects. First histological lesions and cardiac dysfunction parallel those found in the human DMD, both in terms of nature and of timeline (23–27). Indeed, LV global systolic dysfunction (*i.e*., decrease of the LVEF or LVFS) occurs at a late stage, long after the onset of locomotor and respiratory signs. Secondly, a cardiac dysfunction can also be detected very early, using DTI-echocardiography, and strain rate decrease notably (24, 28). Third, myocardial fibrosis can be detected non-invasively using MRI and LGE (26). At last, DMD-like ECG abnormalities were described, including deep Q-waves, sinus tachycardia, shortened PR, and premature ventricular contractions (25, 29). However, no ECG longitudinal study was performed on the long-term and using Holter ambulatory recordings, and correlation with cardiac function data are lacking. Furthermore, no HRV assessment was ever performed in GRMD dogs.

This study, based on longitudinal Holter ECG data analysis, including HRV assessment, aimed at describing the sequential events of cardiac involvement in this model to improve its knowledge and optimize its use in preclinical studies. It also aimed at assessing possible correlations of ECG abnormalities with echocardiography and biomarker data, which could be translationally relevant, to improve the knowledge on the DMD cardiomyopathy.

## Material and methods

### Animals

All procedures were performed in accordance with EU Directive 2010/63/EU for animal experiments, and were approved by the Ethical committee of EnvA, ANSES and UPEC under the approval numbers 20/12/12-18 and 2020-02-04-07 (APAFiS nbr #23763-2019112814263892 v7).

Fifteen GRMD dogs were included at the age of two months after genotyping as previously described (44). The recruitment spanned over three years (march 2014 to march 2017), and the dogs were then followed-up their whole life long. When necessary, the GRMD dogs were treated for complications of the disease (mainly antibiotics in case of pneumonia, gastrostomy tube feeding in case of pronounced dysphagia), but no steroid nor other known interfering drug (*e.g.*, ACE inhibitors, β-blockers) were prescribed. Nine healthy littermates housed and cared in the same conditions, recruited from March 2014 to August 2020, were used as controls, and were followed-up until either 6 months of age (n=3) or 24 months of age (n=6), before being rehomed. All these dogs were part of a multi-parametric natural history study.

### Holter ECG-recordings

Ambulatory six-lead Holter ECG were recorded overnight in normal housing conditions using a telemetry device (EmkaPACK 4G, Emka technologies, Paris, France), allowing for 12 hour-recordings. Electrodes were clipped on four skin adhesive patches placed at standardized positions after shaving: LA electrode was positioned on the left at the level of the precordial shock, and RA symmetrically on the left side; RL and LL electrodes were respectively positioned in the alignment of RA and LA, at the level of the umbilicus. Electrodes wires were connected to an emitter, which was placed in a dedicated jacket worn by freely moving dogs. ECG was acquired at a sampling rate of 500 Hz using the software IOX (Emka technologies).

Iterative recordings were performed at the following ages: 2, 4, 6, 9, 12, 18, 24 months of age. In surviving GRMD dogs, recordings were renewed at 36, 48 and 60 months of age. A recording was also performed at time of cardiac decompensation for 3 of the GRMD dogs (respectively at 63, 65 and 78 months of age). The number of dogs recorded at each age is available in Tables 1 and S1. Figures in the main paper represent the data from the 2 to 24 months period. The values of GRMD dogs measured at later timepoints are illustrated in figure S4.

**Table 1.**
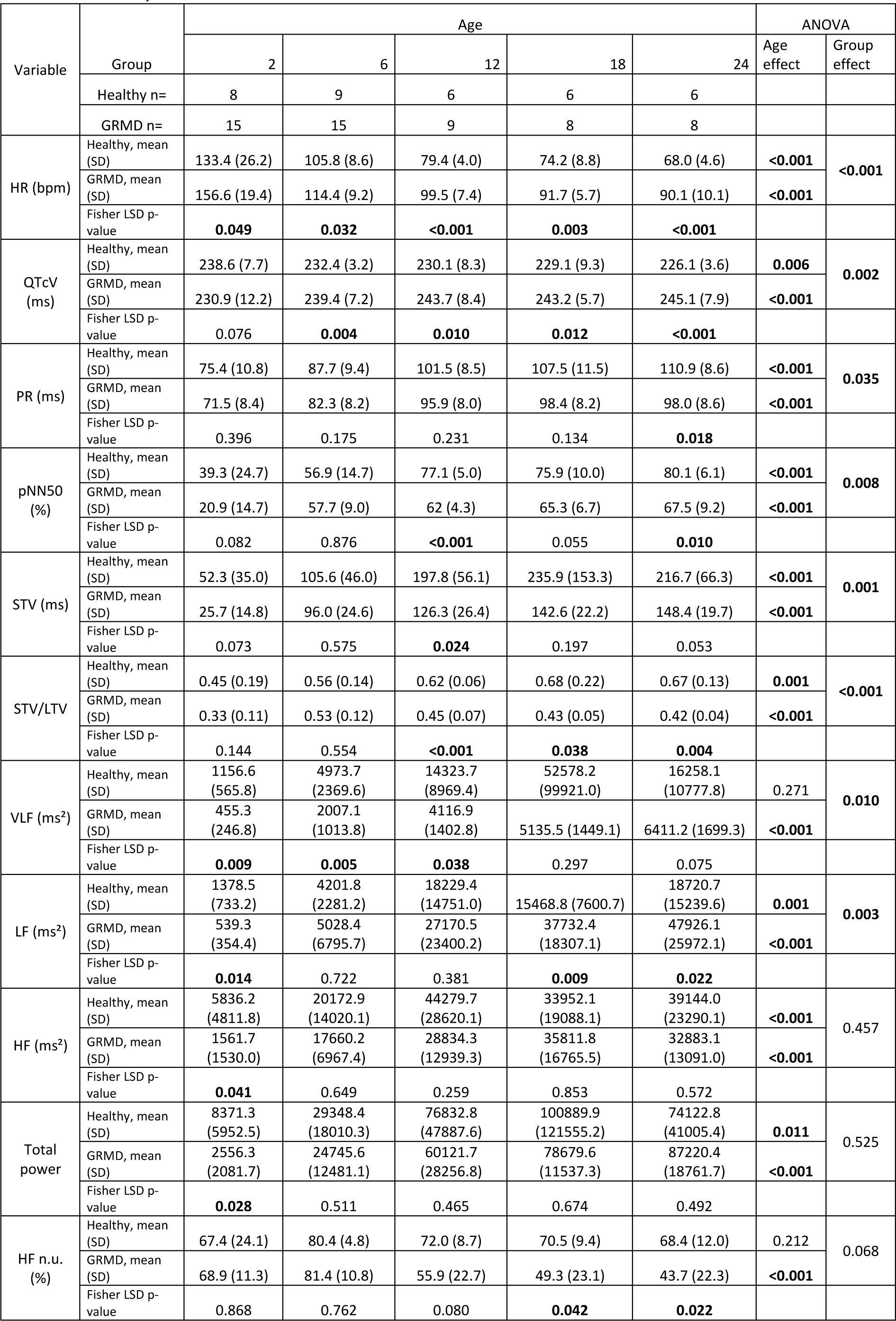
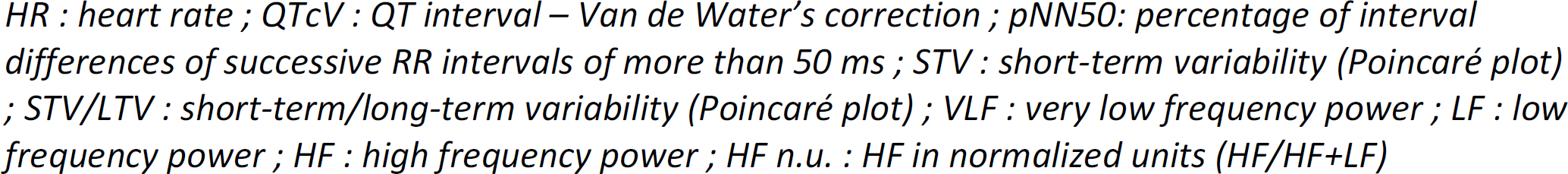
Summary of the main results.

| Variable | Group | Age |  |  |  |  | ANOVA |  |
| --- | --- | --- | --- | --- | --- | --- | --- | --- |
|  |  | 2 | 6 | 12 | 18 | 24 | Age effect | Group effect |
|  | Healthy n= | 8 | 9 | 6 | 6 | 6 |  |  |
|  | GRMD n= | 15 | 15 | 9 | 8 | 8 |  |  |
| HR (bpm) | Healthy, mean (SD) | 133.4 (26.2) | 105.8 (8.6) | 79.4 (4.0) | 74.2 (8.8) | 68.0 (4.6) | **<0.001** | **<0.001** |
|  | GRMD, mean (SD) | 156.6 (19.4) | 114.4 (9.2) | 99.5 (7.4) | 91.7 (5.7) | 90.1 (10.1) | **<0.001** |  |
|  | Fisher LSD p-value | **0.049** | **0.032** | **<0.001** | **0.003** | **<0.001** |  |  |
| QTcV (ms) | Healthy, mean (SD) | 238.6 (7.7) | 232.4 (3.2) | 230.1 (8.3) | 229.1 (9.3) | 226.1 (3.6) | **0.006** | **0.002** |
|  | GRMD, mean (SD) | 230.9 (12.2) | 239.4 (7.2) | 243.7 (8.4) | 243.2 (5.7) | 245.1 (7.9) | **<0.001** |  |
|  | Fisher LSD p-value | 0.076 | **0.004** | **0.010** | **0.012** | **<0.001** |  |  |
| PR (ms) | Healthy, mean (SD) | 75.4 (10.8) | 87.7 (9.4) | 101.5 (8.5) | 107.5 (11.5) | 110.9 (8.6) | **<0.001** | **0.035** |
|  | GRMD, mean (SD) | 71.5 (8.4) | 82.3 (8.2) | 95.9 (8.0) | 98.4 (8.2) | 98.0 (8.6) | **<0.001** |  |
|  | Fisher LSD p-value | 0.396 | 0.175 | 0.231 | 0.134 | **0.018** |  |  |
| pNN50 (%) | Healthy, mean (SD) | 39.3 (24.7) | 56.9 (14.7) | 77.1 (5.0) | 75.9 (10.0) | 80.1 (6.1) | **<0.001** | **0.008** |
|  | GRMD, mean (SD) | 20.9 (14.7) | 57.7 (9.0) | 62 (4.3) | 65.3 (6.7) | 67.5 (9.2) | **<0.001** |  |
|  | Fisher LSD p-value | 0.082 | 0.876 | **<0.001** | 0.055 | **0.010** |  |  |
| STV (ms) | Healthy, mean (SD) | 52.3 (35.0) | 105.6 (46.0) | 197.8 (56.1) | 235.9 (153.3) | 216.7 (66.3) | **<0.001** | **0.001** |
|  | GRMD, mean (SD) | 25.7 (14.8) | 96.0 (24.6) | 126.3 (26.4) | 142.6 (22.2) | 148.4 (19.7) | **<0.001** |  |
|  | Fisher LSD p-value | 0.073 | 0.575 | **0.024** | 0.197 | 0.053 |  |  |
| STV/LTV | Healthy, mean (SD) | 0.45 (0.19) | 0.56 (0.14) | 0.62 (0.06) | 0.68 (0.22) | 0.67 (0.13) | **0.001** | **<0.001** |
|  | GRMD, mean (SD) | 0.33 (0.11) | 0.53 (0.12) | 0.45 (0.07) | 0.43 (0.05) | 0.42 (0.04) | **<0.001** |  |
|  | Fisher LSD p-value | 0.144 | 0.554 | **<0.001** | **0.038** | **0.004** |  |  |
| VLF (ms<sup>2</sup>) | Healthy, mean (SD) | 1156.6 (565.8) | 4973.7 (2369.6) | 14323.7 (8969.4) | 52578.2 (99921.0) | 16258.1 (10777.8) | 0.271 | **0.010** |
|  | GRMD, mean (SD) | 455.3 (246.8) | 2007.1 (1013.8) | 4116.9 (1402.8) | 5135.5 (1449.1) | 6411.2 (1699.3) | **<0.001** |  |
|  | Fisher LSD p-value | **0.009** | **0.005** | **0.038** | 0.297 | 0.075 |  |  |
| LF (ms<sup>2</sup>) | Healthy, mean (SD) | 1378.5 (733.2) | 4201.8 (2281.2) | 18229.4 (14751.0) | 15468.8 (7600.7) | 18720.7 (15239.6) | **0.001** | **0.003** |
|  | GRMD, mean (SD) | 539.3 (354.4) | 5028.4 (6795.7) | 27170.5 (23400.2) | 37732.4 (18307.1) | 47926.1 (25972.1) | **<0.001** |  |
|  | Fisher LSD p-value | **0.014** | 0.722 | 0.381 | **0.009** | **0.022** |  |  |
| HF (ms<sup>2</sup>) | Healthy, mean (SD) | 5836.2 (4811.8) | 20172.9 (14020.1) | 44279.7 (28620.1) | 33952.1 (19088.1) | 39144.0 (23290.1) | **<0.001** | 0.457 |
|  | GRMD, mean (SD) | 1561.7 (1530.0) | 17660.2 (6967.4) | 28834.3 (12939.3) | 35811.8 (16765.5) | 32883.1 (13091.0) | **<0.001** |  |
|  | Fisher LSD p-value | **0.041** | 0.649 | 0.259 | 0.853 | 0.572 |  |  |
| Total power | Healthy, mean (SD) | 8371.3 (5952.5) | 29348.4 (18010.3) | 76832.8 (47887.6) | 100889.9 (121555.2) | 74122.8 (41005.4) | **0.011** | 0.525 |
|  | GRMD, mean (SD) | 2556.3 (2081.7) | 24745.6 (12481.1) | 60121.7 (28256.8) | 78679.6 (11537.3) | 87220.4 (18761.7) | **<0.001** |  |
|  | Fisher LSD p-value | **0.028** | 0.511 | 0.465 | 0.674 | 0.492 |  |  |
| HF n.u. (%) | Healthy, mean (SD) | 67.4 (24.1) | 80.4 (4.8) | 72.0 (8.7) | 70.5 (9.4) | 68.4 (12.0) | 0.212 | 0.068 |
|  | GRMD, mean (SD) | 68.9 (11.3) | 81.4 (10.8) | 55.9 (22.7) | 49.3 (23.1) | 43.7 (22.3) | **<0.001** |  |
|  | Fisher LSD p-value | 0.868 | 0.762 | 0.080 | **0.042** | **0.022** |  |  |

*HR : heart rate ; QTcV : QT interval – Van de Water’s correction ; pNN50: percentage of interval differences of successive RR intervals of more than 50 ms ; STV : short-term variability (Poincaré plot) ; STV/LTV : short-term/long-term variability (Poincaré plot) ; VLF : very low frequency power ; LF : low frequency power ; HF : high frequency power ; HF n.u. : HF in normalized units ( $HF/(HF+LF)$ )*

### ECG analysis

Quantitative ECG analysis was performed in the software ecgAUTO (Emka technologies), on the lead II. For each trace, a waveform library including normal beats and possible arrhythmic events was built manually by scanning each trace. All the analyses were performed by the same operator. Automatic library-based analysis was then run, and the following parameters were calculated and averaged over the whole trace: heart rate (HR (beats/min)), Q/R ratio, PR interval (ms), QT interval (ms). In order to compensate for HR variations among dogs and ages, the best QT correction formula to be applied to this cohort was evaluated in a preliminary study based on the data obtained in the healthy dogs, as recommended in the literature (45). The following corrections were assessed: QTcB, QTcF, QTcV, and QTcM (45). The correction formula which was the best in decorrelating QT values from heart rate was kept for the subsequent analysis.

The arrhythmic events were categorized and their number was quantified and normalized by the total amount of detected beats over the trace. Concerning premature ventricular beats (PVBs), the number of doublets, triplets or salvos was quantified, and each trace was stratified according to the Lown classification of PVBs severity (0: no PVB; 1: rare isolated PVBs <30/hr; 2: frequent isolated PVBs >30/hr; 3a: isolated polymorphic PVBs; 3b : bigeminated PVBs; 4a: doublets or triplets; 4b: ventricular tachycardia (> 3 consecutive PVBs); 5: R/T phenomenon) (46).

The presence or absence of fragmented QRS was assessed visually on the traces.

### Heart rate variability (HRV) analysis

The HRV analysis was performed using the software ecgAUTO (Emka technologies) after exclusion of the arrhythmic beats and the normal beats preceding and following each arrhythmic event.

Time-domain analysis was first performed providing classical HRV indices, including SDNN (standard deviation of the RR intervals), CV(RR) (coefficient of variation of the RR intervals, *ie* the SDNN/meanRR ratio), RMSSD (square root of the mean squared differences of successive RR intervals), HRV triangular index (total number of RR divided by the number of RR on the highest bin of the RR-density histogram constructed after a 128 Hz sampling), and pNN50 (percentage of interval differences of successive RR intervals of more than 50 ms) (22). A pNN10%RR was also calculated as the percentage of interval differences of successive RR intervals of more than 10% of the mean RR on the trace, to adapt to differences in HR between ECGs, groups and ages.

Poincaré plot analysis was performed by plotting each RR to the previous one. Qualitative visual analyses of the plots were performed, and associated with a quantitative analysis: short-term and long-term variability (STV and LTV (also commonly called SD1 and SD2) were calculated. LTV was the standard deviation of the points along the RR=RR-1 line (line of identity), and STV was the standard deviation of the points along a perpendicular crossing this line of identity at the mean RR point value (30).

Frequency-domain analysis of the RR intervals was performed after 512-points Fast Fourier Transform on 15-minute-long epochs and a rectangular windowing, allowing the calculation of the Very Low, Low and High Frequency powers (VLF, LF and HF) respectively corresponding to the areas under the curve of the spectrum within the following frequency bands: 0.003-0.04 Hz, 0.04-0.15 Hz and 0.15 Hz-0.4 Hz respectively. The mean LF/HF ratio across the epochs, and the mean HF n.u. (normalized units, HF n.u.= HF/(LF+HF)) were calculated and used in the analysis (22).

### Echocardiography and serum biomarkers

The dogs included in this study were part of a larger natural history study and also participated in a study on echocardiography in GRMD dogs, recently published (24). Echocardiography was performed at the same timepoints as the ECG recordings (*i.e.*, 2, 4, 6, 9, 12, 18 and 24 months of age, and every year thereafter for GRMD dogs), as previously described (24). A specific focus on two echo indices was made in the present study: the endo-epicardial gradient of velocity, which is impaired very early in GRMD dogs, and the left ventricular fractional shortening (LVFS) that reveals the occurrence of dilated cardiomyopathy in the adulthood of these dogs. Blood samples were collected from the dogs at the same timepoints and cTpnI and NT-proBNP (biomarkers of myocardial damage and stress, respectively) were measured.

### Statistical analysis

The best QT correction formula was assessed by calculating a Pearson coefficient of correlation between QTc and HR, and by determining the slope of the regression line.

A repeated measures ANOVA was used to assess the age effect (within effect) in each group (GRMD, Healthy). The overall group (GRMD vs healthy) effect was assess using a mixed effects model followed by a Fisher’ LSD test to assess group effect at each time point.

Correlation between the indices demonstrated to be different in GRMD dogs relative to healthy dogs, and LVFS at the age of 24 months was assessed in the 8 surviving GRMD dogs, using a Spearman coefficient of correlation. The level of statistical significance was set at *p* ≤ 0.05.

## Results

### GRMD dogs

Fifteen GRMD dogs were included in this study at the age of 2 months, and underwent iterative Holter ECGs. Four of them were euthanized around six months of age due to a loss of ambulation. Two dogs were euthanized around 9 months of age. Nine dogs were still alive at 12 months of age. A seventh dog was euthanized at the age of 15 months. Eight GRMD dogs were still alive at the 18 and 24 months timepoints. Three dogs died in their 3^rd^ year. One died at 36 months of age around one hour after uncomplicated recovery from general anesthesia, after a short syncopal episode followed by cardiorespiratory arrest and failed resuscitation attempt. The four last GRMD dogs all survived until the 60 months timepoint. Two of them decompensated their cardiac insufficiency at respectively 63 and 65 months of age. A third one had to be euthanized at 70 months of age due to idiopathic pericardial effusion. The last surviving GRMD dog of this cohort decompensated his cardiac insufficiency at 78 months of age. A survival curve is available in the Figure S1 together with a more detailed description of the causes of death of the animals. The number of GRMD dogs per age and the main results are available in Tables 1 and S1

### ECG analysis

**Heat rate** (HR) significantly decreased with age in both healthy and GRMD dogs and was significantly higher in GRMD dogs than healthy dogs over the 2-24 months period (*p* < 0.001). GRMD dogs exhibited significantly higher HR relative to healthy littermates at 2 months of age, and from the age of 6 months. From the age of 12 months, the means of the HR in the two groups basically differed from 20 bpm. A marked increase in HR was observed in the three dogs with decompensated cardiomyopathy. (Figure 1A)

**Figure 1:**
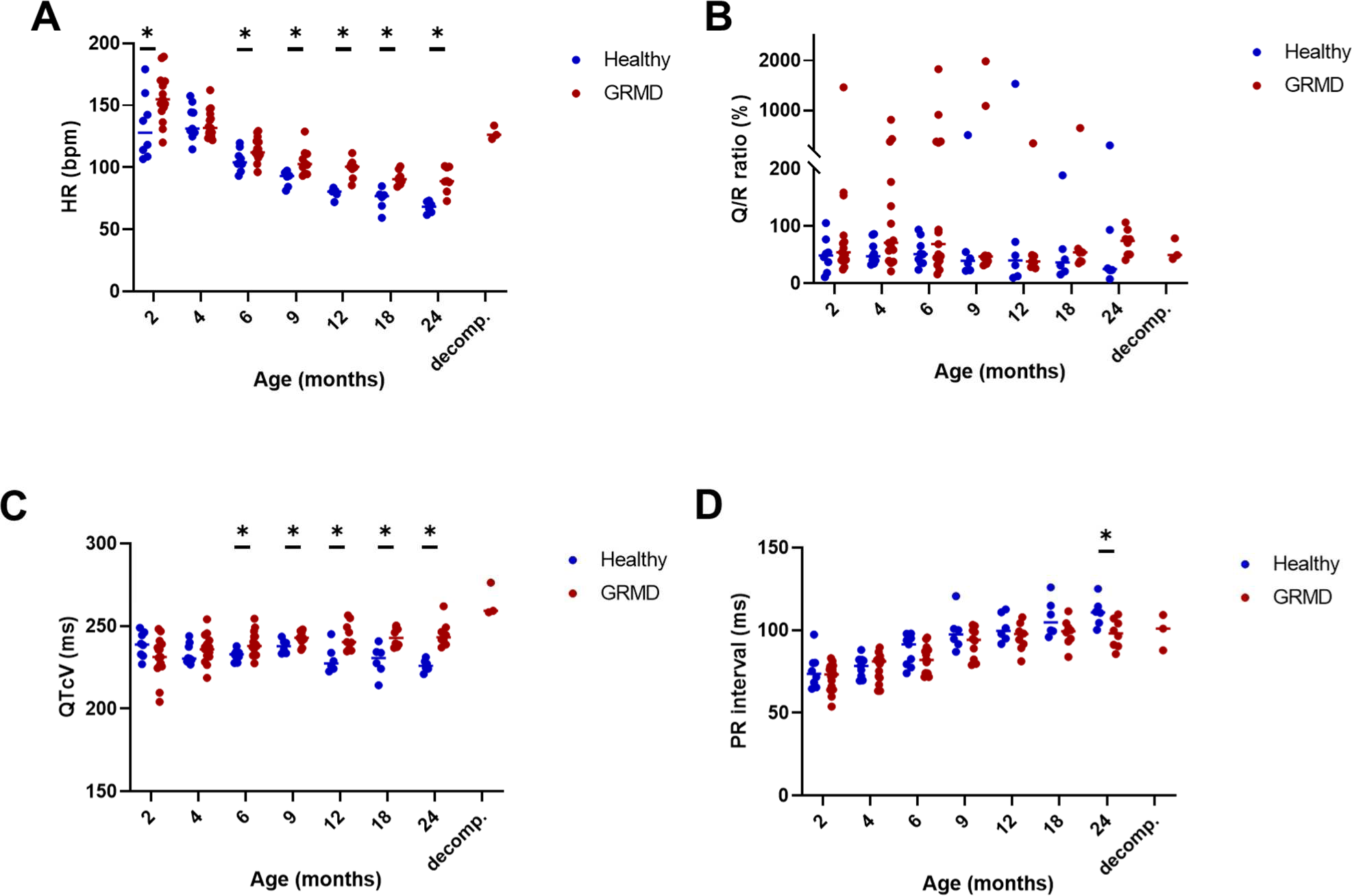
ECG analysis – evolution of intervals and waves ratio with age. On all the graphs, the evolution of GRMD dogs’ values are represented by red dots, and healthy dogs’ values by blue dots. A line indicates the median of each population at each timepoint. Asterisks symbolize a significant difference (Fisher LSD test p<0.05) between GRMD and healthy dogs at a given timepoint. **A.** Heart rate (HR) decreased with age in both groups, but was higher in GRMD dogs than in healthy dogs. Significant differences were found at each timepoint except at 4 months of age. HR re-increased in dogs with cardiac decompensation. **B.** The Q/R ratio increased in some individuals, markedly for some of them. No significant difference with healthy dogs was found. **C.** The QTcV (Van de Water’s correction) interval was prolonged in GRMD dogs relative to healthy dogs, with a significant difference from the age of 6 months. QTcV was even more prolonged at the stage of decompensation. **D.** The PR interval increased with age in both groups, but was shorter in GRMD dogs than in healthy dogs at the latest timepoints. The difference was significant at the age of 24 months.

**The Q/R waves amplitude ratio** was calculated to detect possible deep Q-waves in GRMD dogs as previously described (25, 29). This ratio was not found to vary with age. No significant difference between healthy and GRMD dogs could be demonstrated at any age. However, remarkably increased values were observed in some dogs, more frequently in GRMD dogs. It is noteworthy that, including in healthy dogs, on a given trace, Q waves could be found deep at some periods and normal at some other periods suggesting possible position-related change of morphology. (Figure 1B)

**The QT interval** significantly increased with age in both groups, consistently with the HR decrease (Figure S2). No significant difference was found between healthy and GRMD dogs. Given the significant variations in HR with age and between groups, and the known impact of HR on the QT interval, the use of a corrected QT (QTc) appeared necessary to compare both groups. A preliminary study intending to determine the best QT correction formula for our cohort was performed, and based on the HR and four different QTc values obtained in the healthy dogs (Figure S2). A significant negative correlation between QT and HR was confirmed (R = -0.929, slope of the regression line -0.59). As also described in dogs, QTcB, the correction formula used in most of the studies in humans, was found to strongly overcorrect QT values (R = 0.881, slope= 0.53) (28). The QTc with the R and slope values of the regression line closest to 0 was QTcV (R = 0.171, slope of the regression line = 0.04). QTcV was thus selected for the analysis.

A significant increase of QTcV with age was found both in healthy and GRMD dogs and QTcV was significantly increased in GRMD dogs compared to healthy dogs over the 2-24 months period (*p* = 0.002). A significant increase of QTc values was demonstrated in GRMD dogs relative to healthy dogs first at the age of 6 months (*p* = 0.004), and at every timepoint thereafter. The three dogs with decompensated cardiomyopathy exhibited markedly increased QTcV values compared to their values at 60 months of age (Figure 1C, Figure S4).

**The PR interval**, known to be shortened in DMD, was found significantly smaller in GRMD dogs over the 2-24 months period (p = 0.035), and significantly increased with age in both groups (*p* < 0.001) (Figure 1D). The PR interval was significantly decreased in GRMD dogs at the age of 24 months relative to healthy dogs (*p* = 0.018), suggesting that this interval is shortened in GRMD dogs, but this is a rather late ECG finding.

**Arrhythmic events** were frequently detected in GRMD ECG, and most of them were of ventricular origin, though supraventricular arrhythmias or conduction defects were found sporadically. Premature ventricular beats (PVBs) were a common finding in GRMD dogs, and their frequency (in percentage of the total detected beats) increased with age (*p* = 0.007). PVBs were absent from ECGs of 2-month-old GRMD dogs but were found at all ages afterwards and concerned 50 % of the GRMD dogs at the age of 9 months. The severity of these PVBs, according to the Lown classification, tended to increase with age. In GRMD dogs, the two highest grades of the Lown classification, corresponding to ventricular tachycardia salvos, and R/T phenomenon were reached for some GRMD dogs at several timepoints and this could explain the sudden deaths with no significant findings at necropsy sporadically occurring in GRMD dogs (personal communication). In this cohort it was the case for one dog that had R/T PVBs at the age of 4 months and suddenly died, without any identified cause, few days after the 36 months Holter during which the dog presented some episodes of ventricular tachycardia.

It is noteworthy that in the healthy dog group, three individuals presented with PVBs, which occurred relatively frequently for two of them and could manifest as doublets, triplets and salvos for one of them, leading to high scores on the Lown scale. However, PVBs seen in these healthy dogs were always monormorphic (and conserved the same morphology across the different timepoints) while GRMD dogs tended to exhibit polymorphic PVBs. The VT salvos seen in one healthy dog were not sustained, comprising usually 4-5 contractions, and a maximum of 10, once. (Figure 2)

**Figure 2:**
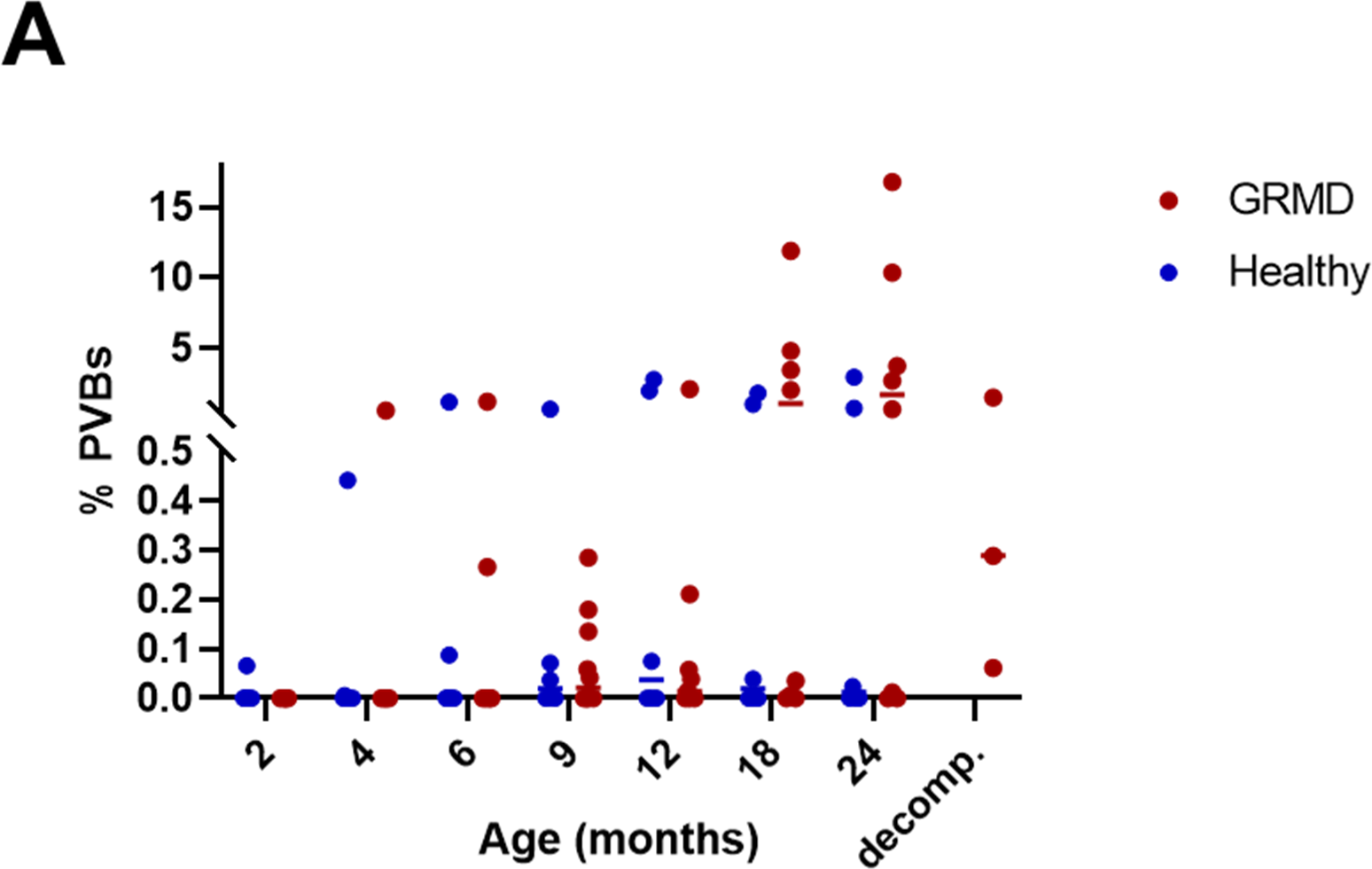
Ventricular arrhythmias in GRMD dogs. **A.** Evolution of the frequency of premature ventricular beats (PVBs, in percentage of the total beats on the trace) with age in GRMD dogs (red dots), versus healthy dogs (blue dots). Lines indicate the median of the population at each timepoint. **B.** Lown classification of PVBs severity shows that the proportion of dogs with PVBs tends to increase with age in GRMD dogs. Severity varies among dogs but can reach high levels with some dogs presenting with ventricular tachycardia or R/T PVBs (about 50% GRMD dogs from the age of 18 months).

**Fragmented QRS** were found in one GRMD dog at 9 months of age, and in four GRMD dogs (half the group) from the age of 24 months. The longest survivor had fragmented QRS detected only at time of decompensation.

### Heart rate variability (HRV) analysis

**Time-domain analysis** parameters all significantly increased with age in both groups, signing an increase in HRV with growing in dogs. No difference was observed between groups for SDNN (standard deviation of the RR intervals) and CV(RR) (coefficient of variation of the RR intervals, *i.e.*, the SDNN/meanRR ratio). A significant decrease in RMSSD (square root of the mean squared differences of successive RR intervals), HRV triangular index (total number of RR divided by the number of RR on the highest bin of the RR-density histogram constructed after a 128 Hz sampling), pNN50 (percentage of interval differences of successive RR intervals of more than 50 ms), and pNN10%(mean RR) (percentage of interval differences of successive RR intervals of more than 10% of the mean RR on the trace) was found in the GRMD group over the 2-24 months period compared to healthy dogs. A significant decrease in pNN50 and pNN10% was found at 9, 12 and 24 months of age, and the decrease was nearly significant at 18 months of age (p = 0.055), as well as at several timepoints from the age of 9 months for the HRV triangular index. For all the time domain analysis parameters, a dramatic drop accompanied the cardiac decompensation. (Figure 3)

**Figure 3:**
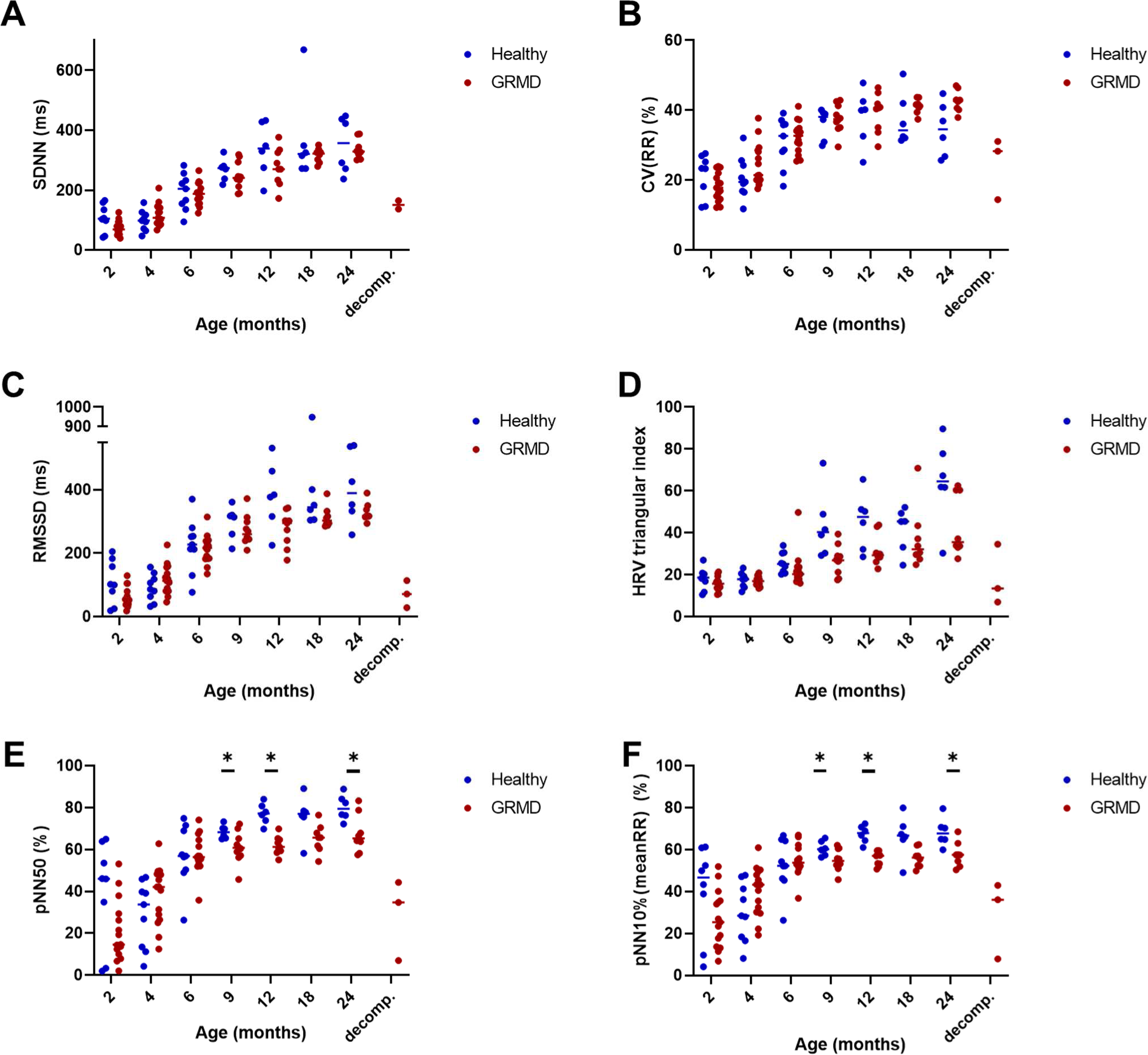
Heart rate variability – Time-domain analysis. On all the graphs, the evolution of GRMD dogs’ values are represented by red dots, and healthy dogs’ values by blue dots. A line indicates the median of each population at each timepoint. Asterisks symbolize significant differences (Fisher LSD test p<0.05) between GRMD and healthy dogs at a given timepoint. **A.** SDNN (standard deviation of the RR intervals), **B.** CV(RR) (coefficient of variation of the RR intervals) and **C.** RMSSD (square root of the mean squared differences of successive RR intervals) all increased with age and with HR decrease in both groups. No difference was found between GRMD and healthy dogs. Cardiac decompensation was associated with a marked decrease of these indices, showing dramatic loss of heart rate variability (HRV). **D.** HRV triangular index tended to be lower in GRMD dogs than in healthy dogs. The difference was nearly significant at 6, 9 and 12 months of age (p= 0.052, 0.053 and 0.057, respectively). **E.** pNN50 (percentage of interval differences of successive RR intervals of more than 50 ms duration) and **F.** pNN10%(meanRR) (percentage of interval differences of successive RR intervals of more than 10% of the mean RR duration) increased with age in both groups. They were found significantly decreased in GRMD dogs at the ages of 9, 12 and 24 months (and nearly significant at 18 months of age, p=0.055). Cardiac decompensation was associated with a marked decrease of both indices.

#### Poincaré plot analysis

The visual assessment of the plots evidenced the typical branched dispersion already described in dogs (30, 31). The longitudinal follow-up also evidenced that this was not seen in puppies but installed with age. In GRMD dogs, the high-density zone close to the line of identity seemed less extended as well as the overall dispersion of the points. Cardiac decompensation in GRMD dogs was associated with a striking modification of the Poincaré plot with a remarkably condensed cloud.

Quantitative analysis was performed by calculating short-term and long-term variabilities (STV and LTV), as well as their ratio. Both STV and LTV significantly increased with age in both groups, and both STV and the STV/LTV ratio were significantly decreased in GRMD dogs compared to healthy dogs over the 2-24 months period. The STV/LTV ratio was found significantly decreased in GRMD dogs from the age of 12 months. Cardiac decompensation was associated with marked STV, LTV and STV/LTV decrease. (Figure 4)

**Figure 4:**
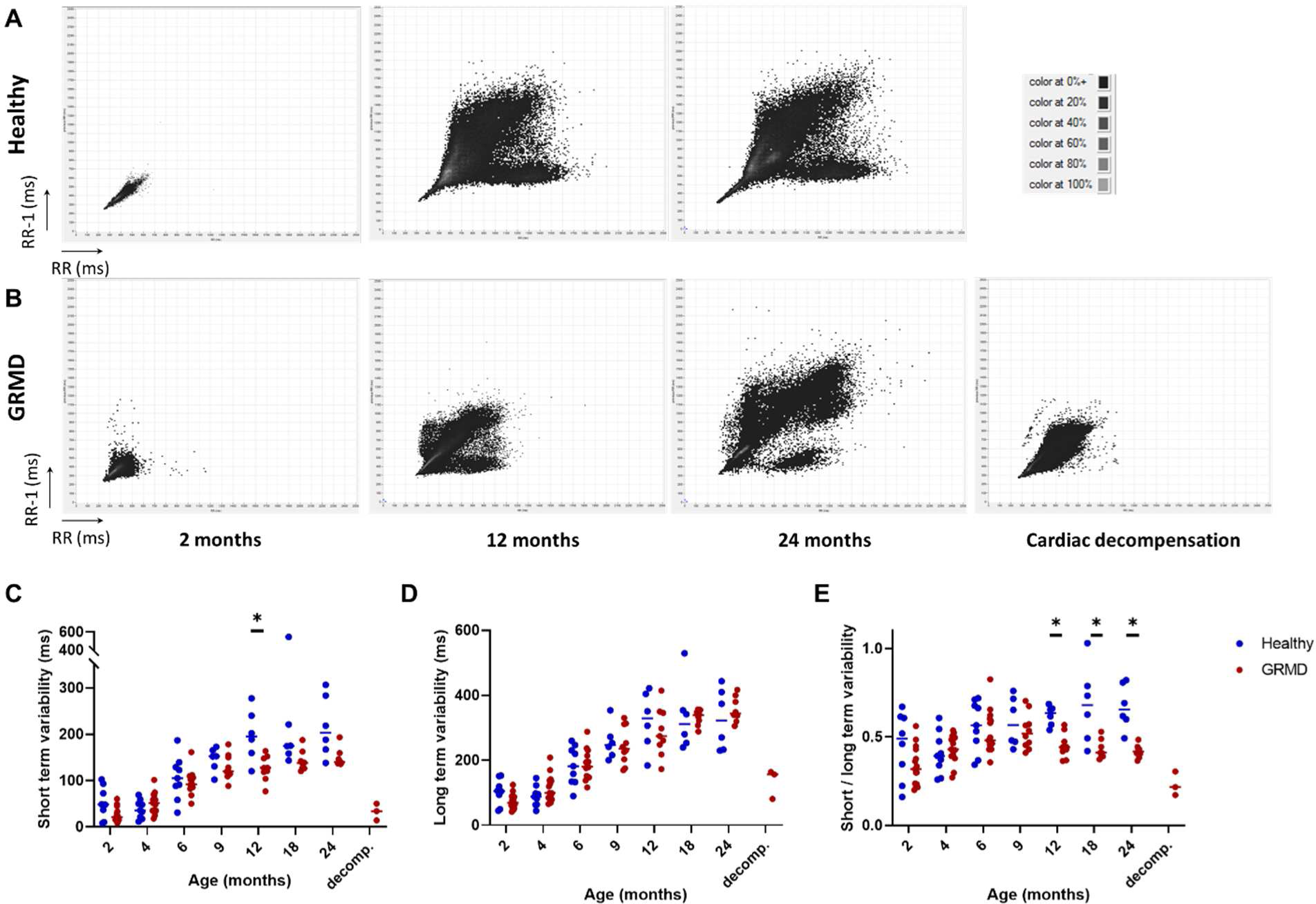
Heart rate variability - Poincaré plot analysis. **A** and **B** show typical evolution of Poincaré plots with age in a healthy and a GRMD dog, respectively. Poincaré plots were constructed by plotting each RR interval to the previous one. The scale was set at 2500 ms x 2500 ms for each graph. Examples at 2, 12, 24 months of age, and at cardiac decompensation for the GRMD dog. The dispersion of the dots on the plot increased with age, and the typical Poincaré plot aspect described in dogs was found at adulthood. The high-density zone was more condensed in GRMD dogs than in healthy dogs and the arms of the Y seemed less dense. At cardiac decompensation in GRMD dogs, this typical Y aspect was completely lost. **C.** Short-term variability (STV) tended to be lower in GRMD dogs, but this was only significant at the age of 12 months (and nearly significant at 24 months of age, p=0.053). **D.** Long-term variability (LTV) did not differ between GRMD and healthy dogs. **E.** The STV/LTV ratio decreased in GRMD dogs relative to healthy dogs, and this difference was significant from the age of 12 months. All three indices dropped at time of decompensation.

#### Frequency-domain analysis

Very low frequency (VLF), low frequency (LF) and high frequency (HF) power components, as well as the total power, increased with age in both groups, and a significant difference between GRMD and healthy dogs was evidenced for VLF and LF over the 2-24 months period. At the age of 2 months, a significant decrease of the total power was found, as well as significant decrease of all three components. Total power was not significantly different between groups thereafter. LF power tended to be higher in GRMD dogs from the age of 9 months, but the difference was only significant starting from 18 months of age. HF power component was not different between groups at any age (except the 2 month-timepoint). Conversely, VLF power component was lower in GRMD dogs than in healthy dogs from the age of 2 months and at most of the timepoints thereafter. The VLF power dramatically decreased in dogs with decompensated cardiomyopathy. It was also the case for the two other power components and the total power. The LF/HF ratio, which is used to measure the sympatho-vagal balance, significantly increased with age in GRMD dogs (*p* < 0.001), but not in healthy dogs. At the age of 4 months, GRMD dogs exhibited significantly lower LF/HF values relative to healthy dogs, but this tendency rapidly inversed because some GRMD dogs exhibited high LF/HF values. From the age of 12 months, this difference was nearly significant (*p* < 0.10). Consistently, HF expressed in normalized units (HF n.u.) was higher than in healthy dogs at 4 months of age (*p* = 0.014), and then decreased to become significantly lower from the age of 18 months. (Figure 5)

**Figure 5:**
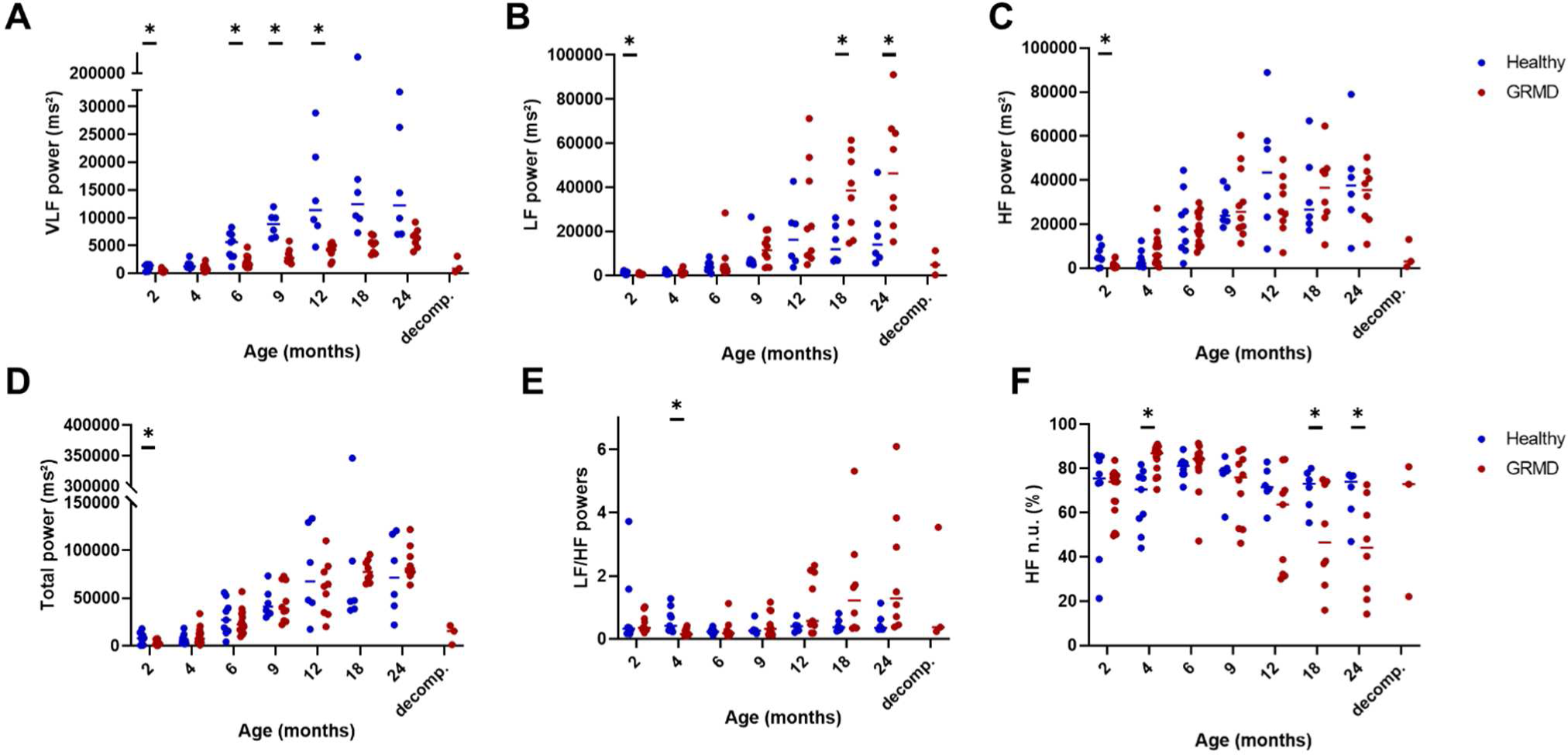
Heart rate variability - Time-frequency analysis. On all the graphs, the evolution of GRMD dogs’ values are represented by red dots, and healthy dogs’ values by blue dots. A line indicates the median of each population at each timepoint. Asterisks symbolize a significant difference (Fisher LSD test p<0.05) between GRMD and healthy dogs at a given timepoint. **A.** Very low frequency (VLF) power was lower in GRMD dogs, and the difference relative to healthy dogs was significant at 2, 6, 9 and 12 months of age. **B.** The low frequency (LF) power tended to increase in GRMD dogs, and the difference relative to healthy dogs was significant at 18 and 24 months of age. **C.** The high frequency (HF) power was unchanged in GRMD dogs relative to healthy dogs. **D.** The total power was not significantly modified in GRMD dogs except at the earliest timepoint where GRMD dogs had significantly lower total power, due to decreased powers in each part of the spectrum. All three power components, as well as the total power markedly decreased in dogs undergoing cardiac decompensation. **E.** Low Frequency / High Frequency (LF/HF) ratio was significantly decreased in GRMD dogs at the age of 4 months, and then increased to some very high individual values in GRMD dogs. The difference between both groups was not significant, but p-values were less than 0.1 from the age of 12 months. **F.** The High frequency power, expressed in normalized units (HF n.u.= HF/(LF+HF)) was consistently increased in GRMD dogs at the age of 4 months, and became decreased thereafter, significantly at the ages of 18 and 24 months.

### Sequence of cardiac changes – comparison with echocardiographic indices and serum biomarkers

The dogs of this study were also part of another study on the natural history of echocardiography in GRMD dogs, recently published (24). This study confirmed that GRMD dogs develop a dilated cardiomyopathy at a late stage of their disease, and that some echocardiography indices can detect an early alteration in LV function. A significant decrease of LVFS relative to healthy dogs was detected at 24 months of age, but the endo-epicardial gradient of velocity was significantly lower in GRMD dogs from the age of 2 months. Some usual cardiac serum biomarkers were also measured in these dogs. The cardiac troponin I (cTpnI) values were significantly increased in GRMD dogs at 4 months of age and tended to be higher thereafter though not reaching statistical significance, showing myocardial damage begins many months before significant LV dysfunction. However, at the individual level, this biomarker appeared to follow a fluctuating evolution with transient peaks and return to lower values. The NT-proBNP values, after a slight increase in some GRMD dogs at 2 months, were within those of healthy dogs until 12 months of age, when some GRMD dogs began to exhibit increased values. The difference relative to healthy dogs became significant at 12 months, to reach very high values in ageing GRMD dogs, which may be due to the progression of myocardial fibrosis (27) (Figure S3).

Taking together the results of previous studies (24, 28) and this study, the sequential evolution of the onset of abnormalities in the occult phase of the dystrophin-deficient cardiomyopathy may thus be drawn. Changes in early echocardiographic indices such as the endo-epicardial gradient of velocity, together with the VLF power component on the ECG spectral analysis and increased HR are the first hallmarks of dystrophin-deficient cardiomyopathy. Then, in the following months, but still preceding the drop of LVFS, QTcV and cTpnI increase, HRV decreases, sympatho-vagal balance modifies, NT-proBNP increases, and arrhythmic events aggravate. Finally, shortened PR and decreased LVFS occur at 24 months of age, indicating the development of dilated cardiomyopathy in young adult dogs.

### Relevance of the ECG indices to LV dysfunction

To understand which parameters could be related to the onset and severity of dilated cardiomyopathy, the correlation between the values obtained for the different ECG and HRV indices proven relevant in GRMD dogs, and their LVFS at the age of 24 months was studied using a Spearman rank test (n= 8 dogs). The endo-epicardial gradient of velocity, as well as the cTpnI and the NT-proBNP values were also included in the analysis. Several significant correlations were found at the age of 4 months. Notably, a negative correlation between HR at 4 months and LVFS at 24 months was found (R = -0.786). A positive correlation between the VLF power at the age of 4 months and the LVFS at 24 months was also found (R = 0.929) (Figure 6). Both HR and VLF were also correlated with the 24 months LVFS at the age of 12 months (R = -0.905 and R = 0.762, respectively). These results suggest that GRMD dogs with elevated HR and low VLF at a young age could develop earlier and/or more severe cardiomyopathy. Probably partly linked to the correlation with HR, pNN10%(meanRR) and pNN50 at 4 months also significantly correlated with LVFS at 24 months (R = 0.762, R = 0.857, respectively) meaning a reduced HRV also could account for a more severe cardiomyopathy. This should however be tempered by the fact that at this timepoint these latter indices were not significantly different between GRMD and healthy dogs. In the same way, a correlation was found between LVFS at 24 months of age and the STV/LTV ratio at 6 months of age (R = -0.738), at a stage when this ratio did not differ between GRMD and healthy dogs. A positive correlation was found between cTpnI values measured at 6 and 18 months of age and the LVFS value at 24 months of age (R = 0.786 and R = 0.714, respectively). This correlation could reflect that a more contractile dystrophin-deficient left ventricle might have elevated risk of cardiomyocyte necrosis. The frequency of PVBs was negatively correlated with the 24 months LVFS both at 18 and 24 months (R = -0.829 and R = -0.732). At last, the PR interval, significantly shorter relative to healthy dogs at the age of 24 months, was significantly correlated with LVFS at this timepoint (R = 0.738). This correlation was also found at 4 and 12 months of age (R = 0.929 and R = 0.762, respectively), however the values were not different from the values of healthy dogs at these earlier timepoints. No correlation was found between the endo-epicardial gradient of velocity and the 24 months LVFS at any timepoint. No correlation was found between NT-proBNP values at any timepoint and 24 months LVFS value. However, these both indices might correlate later on in the disease course: at the age of 36 months, NT-proBNP ranked the GRMD dogs exactly like LVFS did.

**Figure 6:**
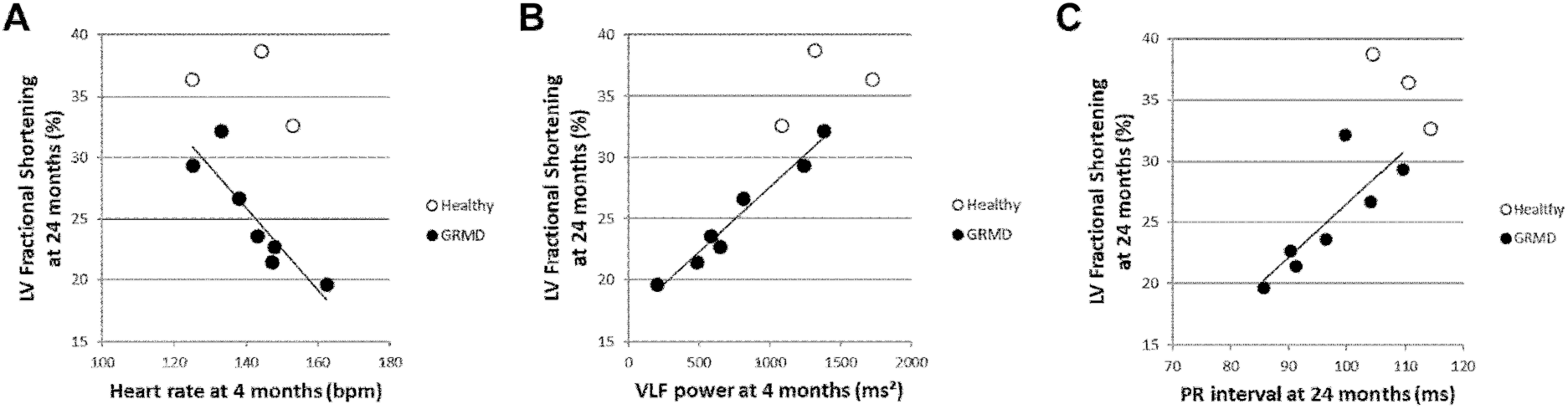
Correlations between ECG indices and fractional shortening at the age of 24 months. Some of the most significant correlations found between ECG indices and the LVFS at 24 months of age in GRMD dogs (black dots) are represented on these graphs; healthy dogs’values are represented by empty dots but were not included in the correlation analysis. **A.** Significant negative correlation between the heart rate at 4 months of age and LVFS at 24 months of age (R=-0.786) **B.** Significant positive correlation between the VLF power at 4 months and LVFS at 24 months of age (R=-0.929) **C.** Significant positive correlation between the PR interval and LVFS at 24 months of age (R=0.738).

## Discussion

This study is the first one to report on longitudinal Holter-ECG findings and to investigate HRV in GRMD dogs. It reveals several similarities with ECG abnormalities found in DMD patients, and reinforces the interest of the GRMD dog as a model for dystrophin-deficient cardiomyopathy. Like in DMD patients, HR was increased, PR interval shortened, QTc prolonged, deep Q waves were found on some dogs, and severe ventricular arrhythmia that could cause sudden death were evidenced (5, 10, 11). Also as described in DMD patients, most of these abnormalities were detected months before the decrease of LVFS. These early hallmarks of the dystrophin-deficient cardiomyopathy can be helpful in evaluating and translating therapeutic effects of preclinical trials performed during the occult phase of the cardiomyopathy.

It is noteworthy that deep Q-waves were only found in some GRMD individuals in our study, and were not correlated with the progression of the disease, whilst it was previously described as a hallmark of ECGs of GRMD dogs (25, 29). It has been reported that deep Q-waves were not found on all DMD patients, and even existing, deep Q-waves do not appear on all leads (10). Since our study only focused on the lead II, it cannot be excluded that deep Q-waves could have been found in more GRMD dogs, with a more interesting evolution pattern, on some other leads. A standardized position, precluded by ambulatory ECG recordings, is also probably important for such analyses of wave sizes, that can vary upon body position (32), and Holter-ECG might thus not be the suitable way to evaluate such abnormalities.

In DMD patients, Holter-ECG is recommended in late non-ambulatory patients to detect severe ventricular arrhythmias (3). Our study shows that ventricular arrhythmias are also commonly found in GRMD dogs, increase with age in terms of severity and correlate with cardiac dysfunction. However, importantly, severe ventricular arrhythmias could be sporadically found in young animals as well. This emphasizes the possible interest to anticipate Holter monitoring in DMD patients.

HRV was found decreased in GRMD dogs. However, the observed modifications were less numerous and striking than reported in DMD patients, using time-domain analysis (15–17, 19, 20). For example, SDNN and RMSSD are decreased in DMD patients, and we did not find them modified in GRMD dogs; pNN50, HRV triangular index and STV/LTV were only modified at a few timepoints. Cardiac decompensation was however associated with striking decrease of all these indices. These few significant modifications before cardiac decompensation could be partly explained by a possible lack of sensitivity of the time-domain indices that were used to assess HRV in dogs. A recent study focused on the differences observed between dogs and humans regarding HRV. This study emphasized on the fact that pronounced respiratory sinus arrhythmia observed in dogs leads to a non-linear and non-gaussian distribution of the RR intervals, dictated by the parasympathetic system influence on the sinus node (31). These differences could account for a possible lack of sensitivity and relevance, in GRMD dogs, of the usual linear indices used to assess HRV. Non-linear indices that also gain interest in humans, should be investigated in future studies (31, 33).

Frequency-domain analysis confirmed that autonomic dysfunction in GRMD dogs resembles the one described in DMD patients. Impaired sympatho-vagal balance was evidenced by an increased LF/HF ratio and a decreased HF n.u., both suggesting a decreased vagal tone and an increased sympathetic activity. This perturbed autonomic balance has been described in young DMD patients, in whom it progresses with age, and has been proposed as a “driving force for myocardial fibrosis” and cardiac function degradation (16, 20). Some studies showed that these perturbations vary with circadian rhythm: in DMD patients, the LF/HF ratio does not decrease at night like in normal patients (15, 16). These circadian variations have not been investigated in our study, but the fact that recordings were performed at night was probably a favorable context to detect the nocturnal LF/HF elevation, progressing with age, in GRMD dogs. With age and disease evolution, some GRMD dogs exhibited very high values of LF/HF ratio and very low values of HF n.u. However, the correlation analysis did not evidence any correlation between the LF/HF ratio or HF n.u. and cardiac function decline.

Interestingly, another component of the power spectrum, the one measured in the very low range of frequencies (VLF), was decreased early in GRMD dogs, and as HR, correlated months before with LVFS values at 24 months of age. This part of the power spectrum has never been deeply investigated in DMD patients to our knowledge, but was reported significantly decreased in one study (14). Our study emphasizes the possible interest to study this part of the spectrum in DMD patients. The mechanisms underlying this VLF component have not been clearly understood, even if it is the major contributor to the total power (21). It has been reported that VLF is related to various extra-cardiac causes, including thermoregulation, renin-angiotensin system, vasomotor tone, and physical activity (34, 35). On this latter point, a comparison of VLF values of GRMD dogs with a loss of ambulation (mean 2732.2 ms² SD 1555.6 ms²) with those still ambulant at the age of 6 months (mean 1743.3 ms² SD 1436.7 ms²) evidenced no difference between both subgroups. At the same timepoint, nocturnal activity measured concomitantly to the ECG, was not significantly different between the GRMD and the healthy groups of dogs. This suggests that decreased physical activity is not the main contributor to the decrease of VLF in GRMD dogs. Core temperature was not measured during ECG recordings but usual rectal temperature measured in GRMD dogs is within normal ranges of the canine species, suggesting thermoregulation is not either a major contributor to the VLF decrease. More recent studies show that VLF is generated intrinsically by the stimulation of afferent sensory neurons in the heart, that in turn activate feedback loops that modulate its frequency and amplitude, towards a decrease by decreasing the parasympathetic outflow and an increase by increasing the sympathetic outflow (21, 36). This new perspective offers VLF an important signification, as a direct reflect of the autonomic nervous system outflow on the intrinsic cardiac nervous system activity (21). VLF was found to be decreased in diverse pathological contexts, probably because it is an integrative marker of autonomic-hormonal control, and it was found to be a powerful independent predictor of severe clinical events (37–41). In the dog model of DMD, we found not only that VLF power was decreased, but also that this decrease was occurring as one of the earliest ECG modifications, with a correlation to the subsequent cardiac dysfunction. This suggests autonomic dysfunction, reflected by decreased VLF (and to a lesser extent by LF/HF and HF n.u.), is an early event in the disease course, and might be a “driving force” for cardiac dysfunction. However, it cannot be excluded that the known vasomotor defect described in GRMD dogs and hypothesized in DMD patients could also account for the VLF decrease and promote cardiac dysfunction (42, 43). If this latter hypothesis would be confirmed, it might support the early use of ACE inhibitors, known to increase VLF, to delay cardiac dysfunction in DMD patients (34, 36).

The endo-epicardial gradient of velocity, measured using Doppler tissue imaging, is also early decreased in GRMD dogs (24). In humans, decreased strain rate also precedes LV global systolic dysfunction (9). However, we did not find any correlation between ECG parameters and subsequent LVFS decrease. This early marker should thus probably be considered as an intrinsic hallmark of dystrophin-deficiency in the heart.

Further studies implicating more dogs should help positioning VLF as a potential early predictor of the cardiac function decline. This study in GRMD dogs, together with data from the literature on the VLF power in other pathological contexts (37–41), should promote studies in DMD patients, intending to investigate the potential interest of the implementation of this marker in the follow-up of patients and to anticipate cardiac dysfunction and adapt therapeutics.

## Acknowledgements

We thank Cyrille Debard in Biovelys (VetAgroSup Lyon) for the Troponin I assessments. We thank the whole team of the Centre d’Elevage du Domaine des Souches – Marshall bioresources for breeding the GRMD dogs.

We thank Lucien Sambin for technical assistance, Dr Pablo Aguilar and the whole team of the CARE-NMD Platform for their loving care on the dogs.

---

**Table S1.**
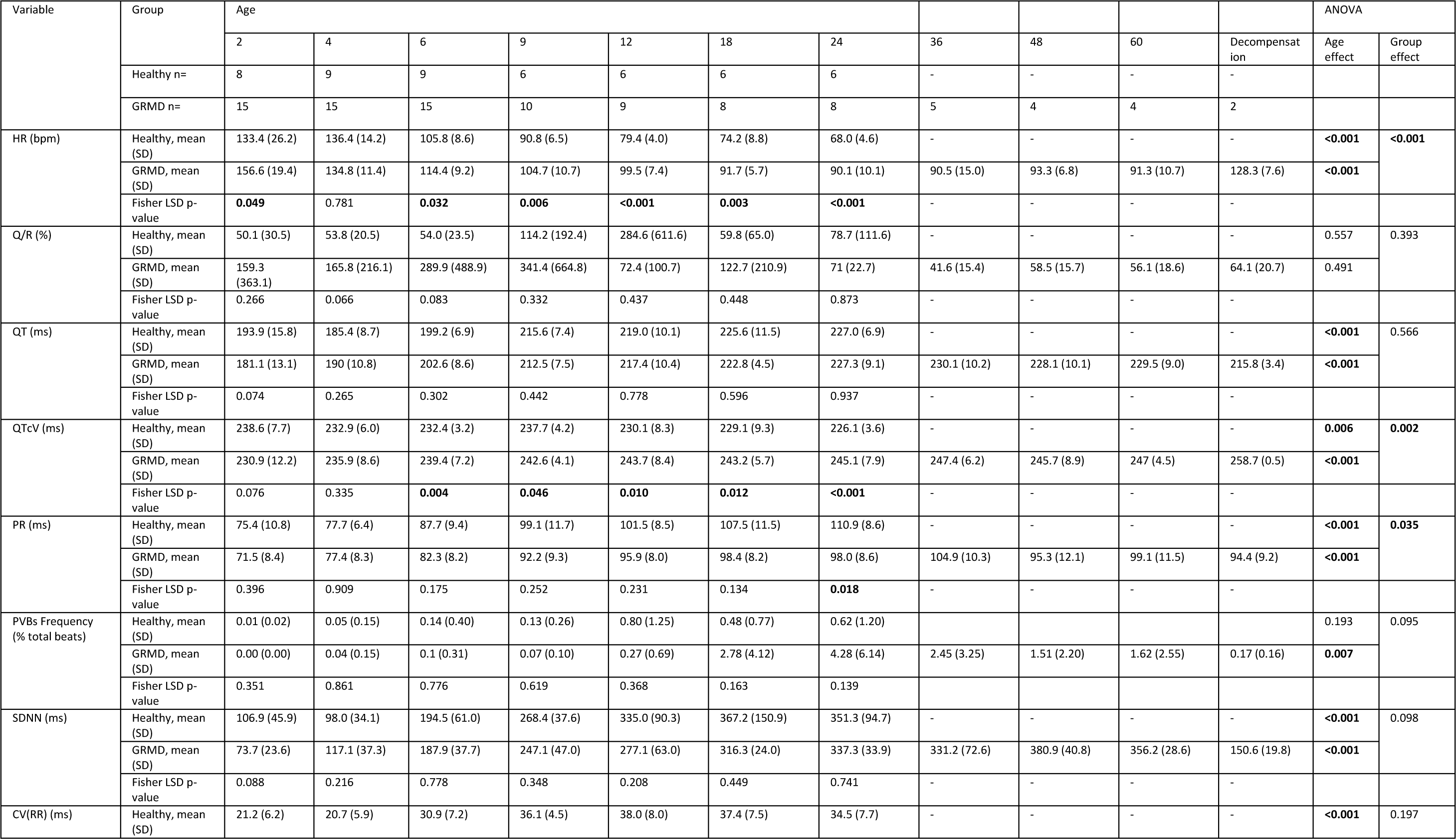
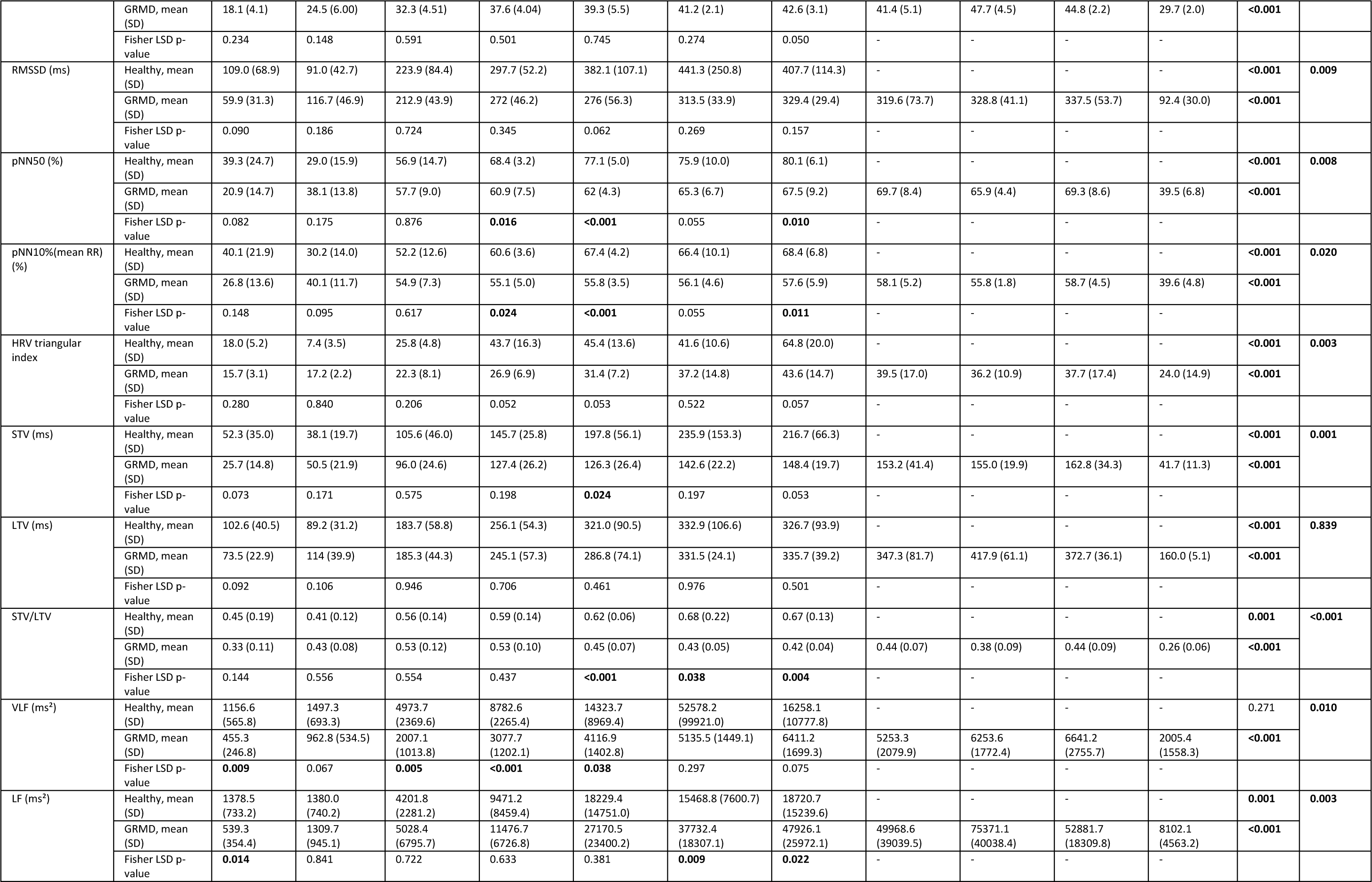
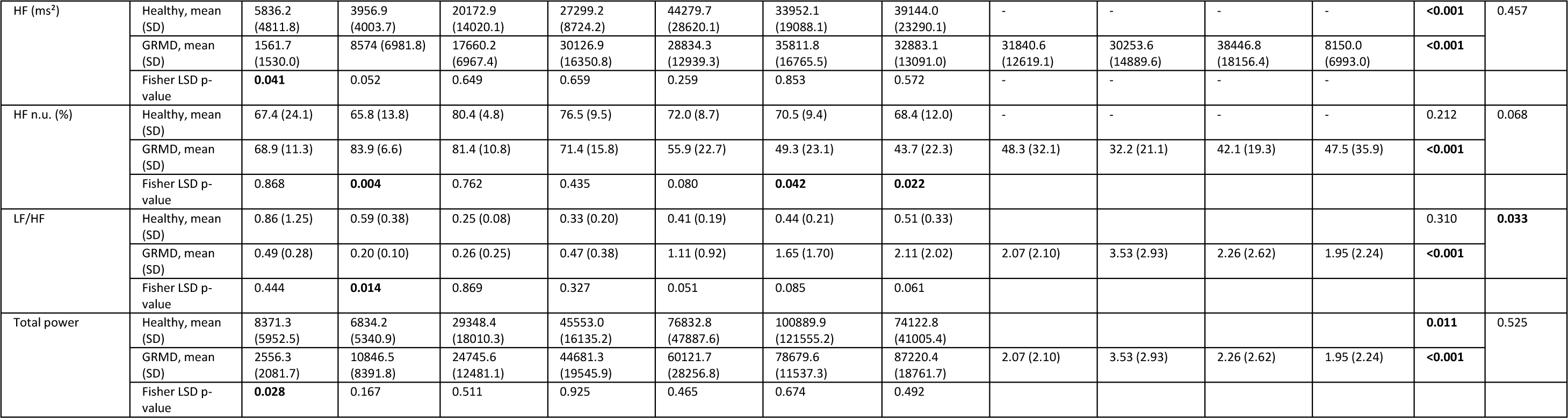
Detailed results.

**Figure S1:**
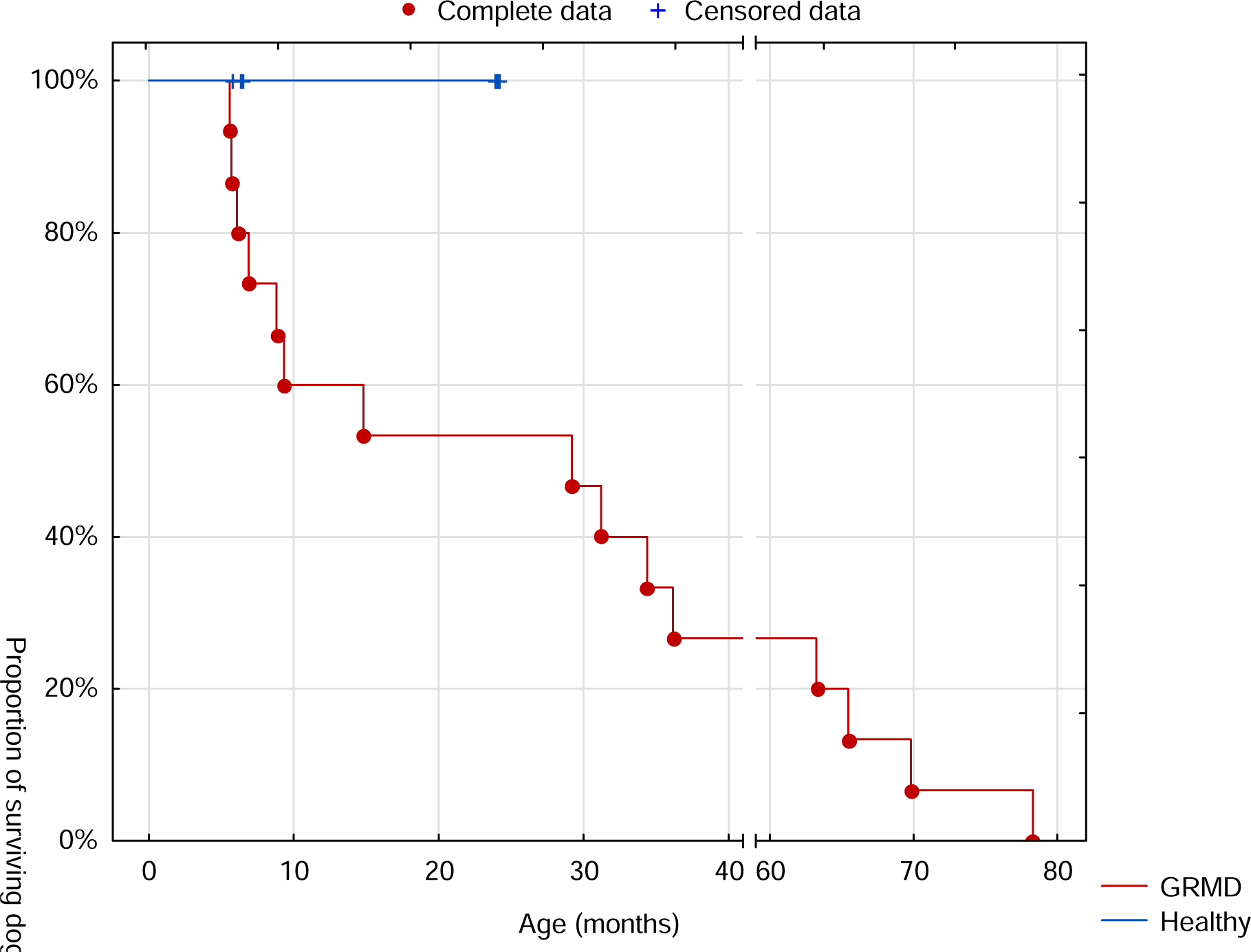
Survival curve of the dogs included in the study.

Nine healthy dogs (blue line) were included and rehomed at 6 (n=3) and 24 months of age (n=6). Fifteen GRMD dogs (black line) were included. Four of them were euthanized around six months of age due to a loss of ambulation and marked dyspnea for one of them. All of them could undergo the 6 months ECG recording. Two dogs were euthanized around 9 months of age, the first one was due to a pulmonary lobar torsion, and the second one in whom the 9 months ECG recording was completed was due to poor mobility. Nine dogs were still alive at 12 months of age. A seventh dog was euthanized at the age of 15 months due to poor mobility associated with marked dyspnea. Eight GRMD dogs were still alive at the 18 and 24 months timepoints. In their third year, two dogs underwent laparotomia intending to reverse an ileus, due in both cases to entangled pylorico-duodenal junction in the hiatal hernia. The proximal part of the descendant duodenum was in both cases necrotic, and the dogs were thus euthanized during surgery. A third GRMD dog died within his third year after a failed resuscitation attempt, following a cardiorespiratory arrest at induction of an anesthesia. Five dogs were still alive at the 36 months timepoint. One of them suddenly died at the age of 36 months around one hour after uncomplicated recovery from general anesthesia during which a short syncopal episode occurred, and the resuscitation attempt failed. The four last GRMD dogs all survived until the 60 months timepoint. Three of them had a decompensated heart failure, manifesting ascites, pleural effusion, and pulmonary edema in one dog, at respectively 63, 65 and 78 months of age and justifying euthanasia. The fourth one, which was in an overall very good shape with a very mild locomotor form of the disease, had to be euthanized at 70 months of age, due to an idiopathic pericardial effusion.

**Figure S2:**
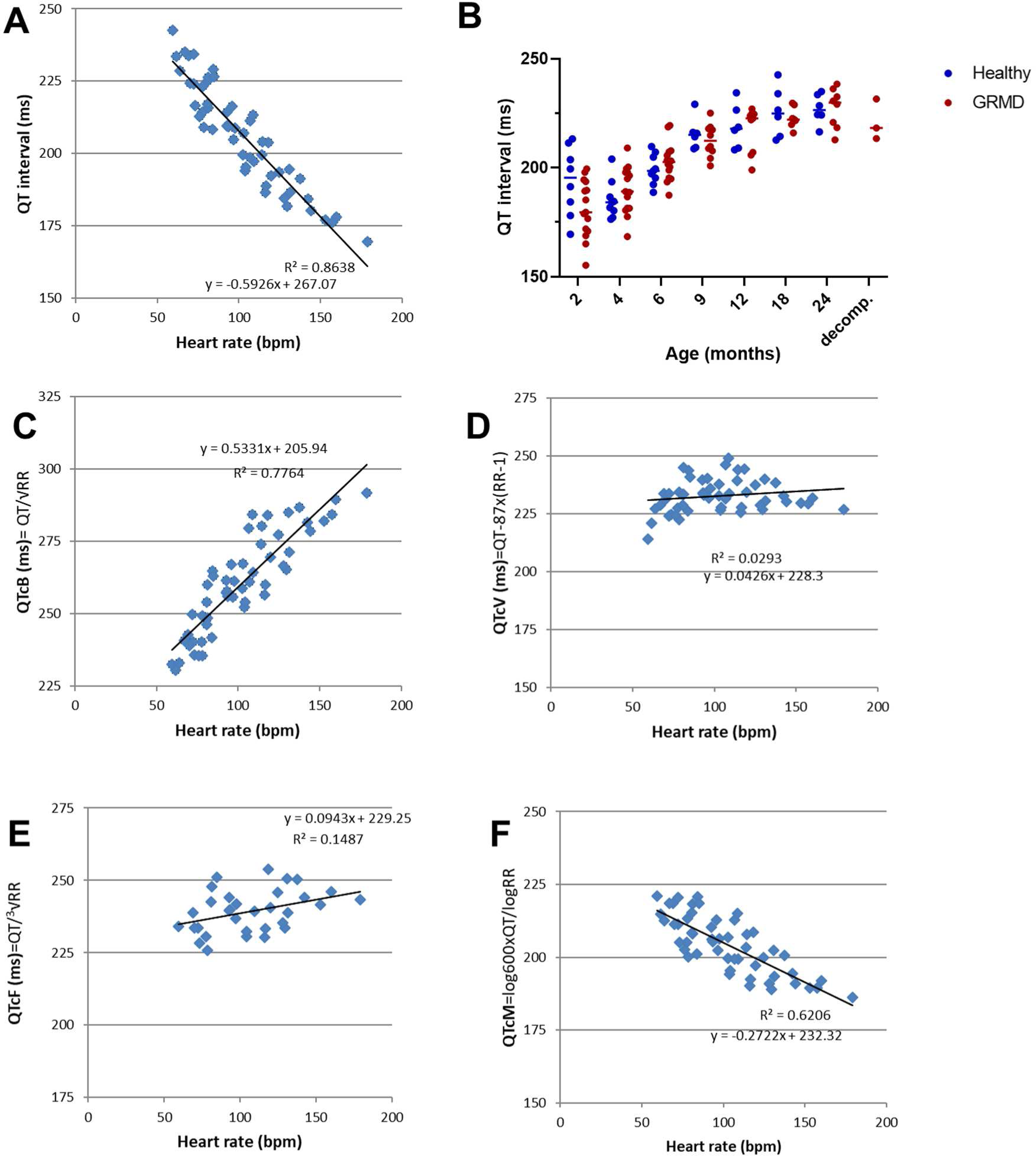
QT correction study.

This study was based on the data obtained on healthy dogs overtime. QTc values according to different formulae (Y axis) were plotted versus HR values (X axis). A: As expected, QT was negatively correlated with HR B: As a consequence, QT increased with age and HR decrease in healthy and GRMD dogs. C. QTcB (Bazett’s correction) overcorrected the QT value, leading to a positive correlation with HR. D. QTcV (Van de Water’s correction) was the best correction formula in decorrelating QT from HR in our dog cohort, consistently with other descriptions in the canine species. E. QTcF (Friedericia’s correction) F. QTcM (Matsunaga’s correction)

**Figure S3:**
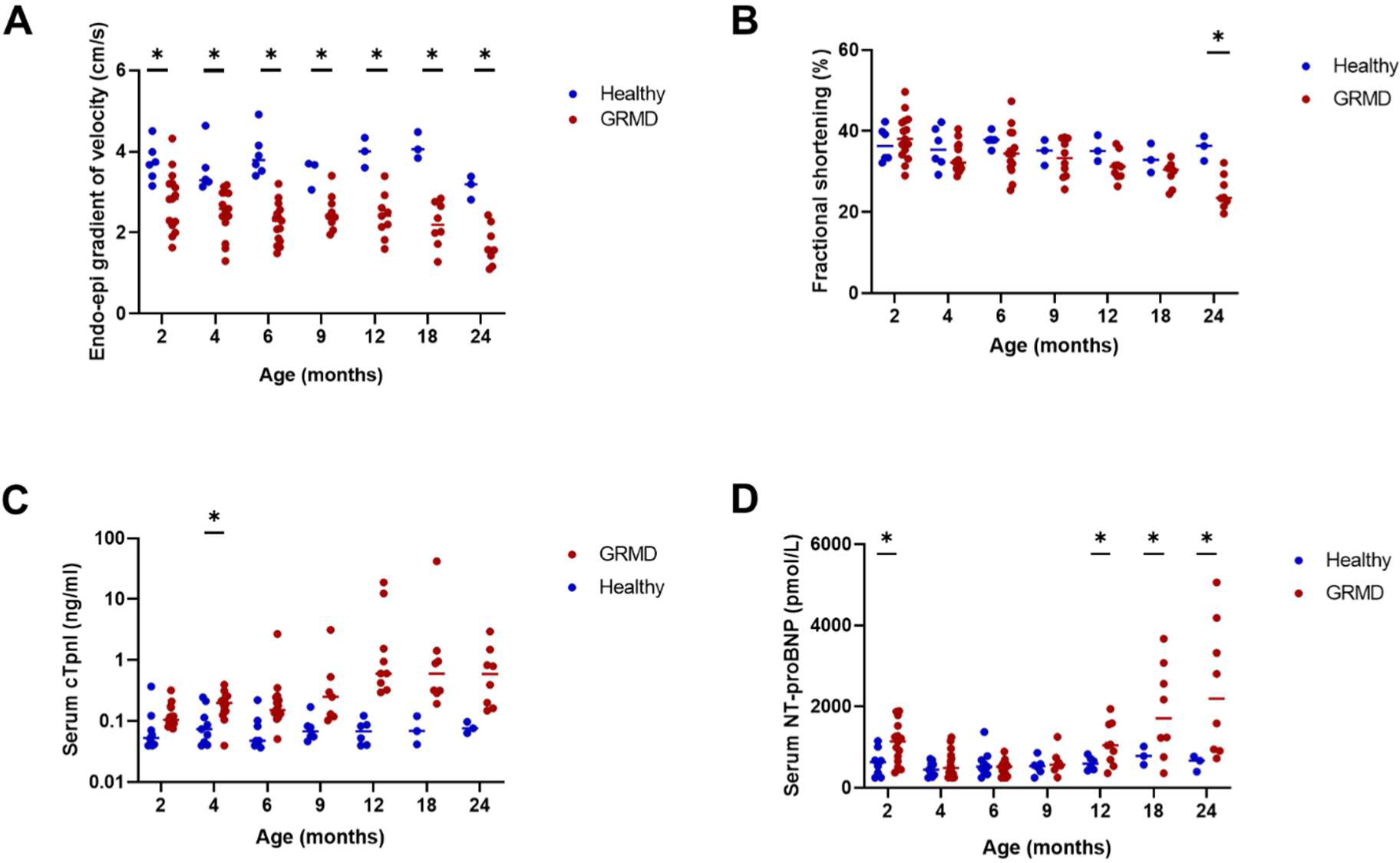
Echocardiographic data and cardiac biomarkers in the studied cohort.

Echocardiography was performed at the same timepoints as Holter-ECG recordings. **A**. Endo-epicardial gradient of systolic velocity was measured using Doppler tissue imaging, and was early decreased in GRMD dogs. It was significantly decreased compared to healthy dogs from the age of 2 months. **B Fractional** shortening was significantly decreased in GRMD dogs at the age of 24 months. Blood samples were also taken at the same timepoints as Holter-ECG recordings. **C.** Serum Troponin I was increased to some very high values attesting to myocardial damage in GRMD dogs (logarithmic scale), though the difference relative to healthy dogs was only significant at the age of 4 months. **D.** NT-proBNP values were slightly increased in some GRMD dogs at 2 months of age, and then remained normal until 12 months when the blood NT-proBNP concentration progressively increased again. NT-proBNP values were significantly increased at the age of 24 months and reached very high values in some GRMD dogs.

**Figure S4:**
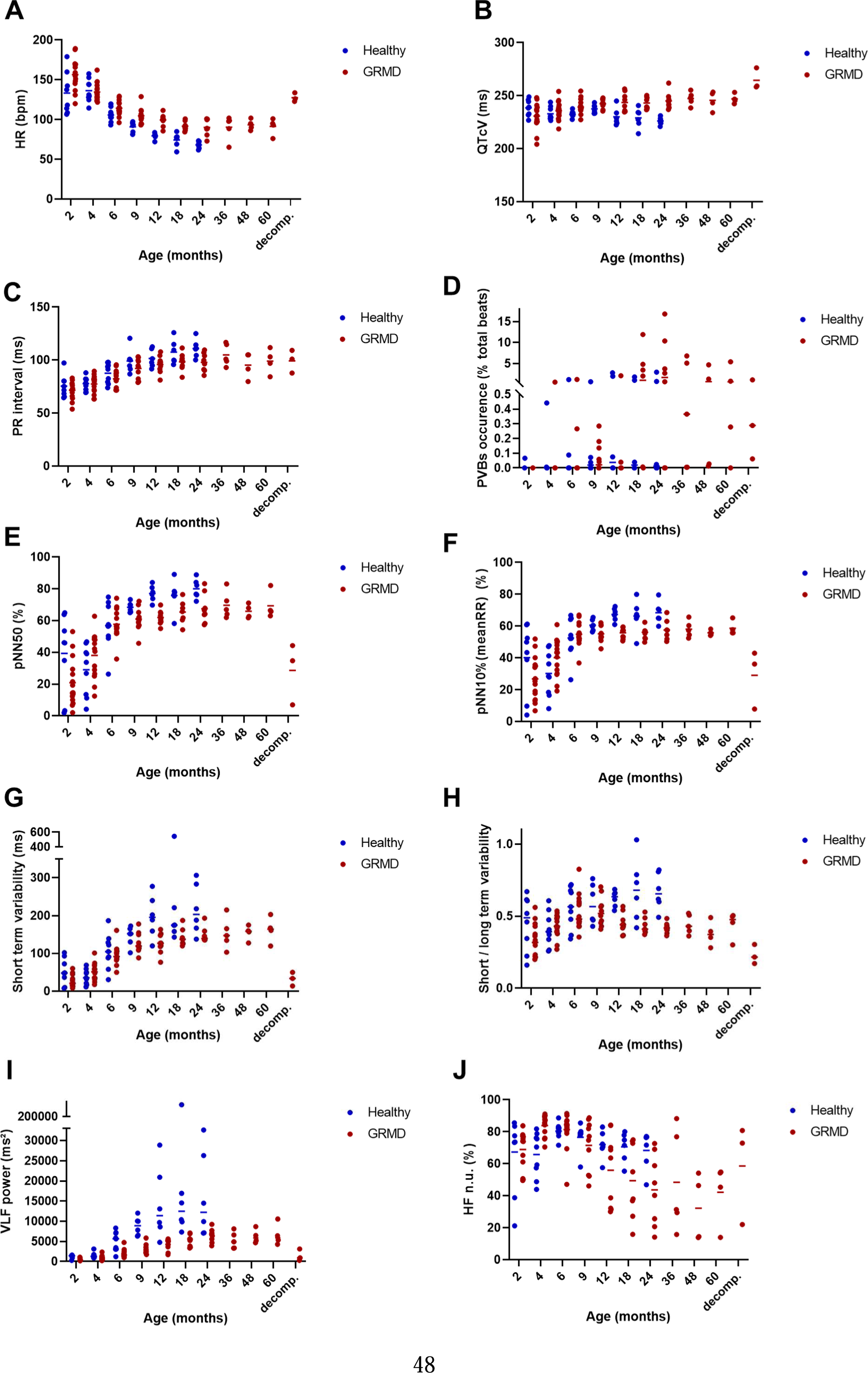
Longitudinal ECG and HRV data including yearly follow-up of GRMD dogs after 24 months of age.

